# Adult sex ratio variation and its sex-specific predictors in shorebirds

**DOI:** 10.1101/2023.08.26.554808

**Authors:** José O. Valdebenito, Tamara Torres-Paris, Juan G. Navedo

## Abstract

The proportion of adult males to females in the adult population, the adult sex ratio (ASR), is an important demographic parameter that has implications in sexual selection, ecology and conservation. ASR variation can be multifactorial but specific variables including sex roles – sex differences in courtship, mate competition, social pair-bonds and parental care– and sex-specific mortality have been suggested as important ASR determinants in birds. However, these relationships have not yet been comprehensively tested in specific avian groups. Here, we used phylogenetic comparative methods to study drivers of ASR variation across shorebirds (Charadrii and Scolopaci; n = 205 species), a charismatic bird group characterised by displaying extreme variations in ecology, sex roles and sexual selection traits. We found that ASR variation is associated with most sex role components but not with their sex-specific mortality. Although sex role and life history variables showed no significant sex bias, we found a trend towards reversed size dimorphism and sex role reversal across shorebirds. Sex roles components also showed correlations among each other that were surprisingly strong and in unexpected directions. Our results confirm that sex roles are important drivers of ASR variation and suggest that shorebirds may have alternative means of sex-specific mortality, possibly linked to their ecology.

## Introduction

The proportion of adult males to females in the adult population –termed adult sex ratio– is a key demographic parameter in natural populations. It has implications in life-history processes such as mate competition and parental care, as well as in conservation because the number of females impacts on population growth (Donald 2007; Hays et al. 2017). In nature, adult sex ratios (ASR) show great diversity and can vary substantially between species and even between populations. For instance, copepods (Cl. Copepoda) are known for having ASRs with strong female biases (Gusmão and McKinnon 2009), while some populations of American red squirrel (*Tamiasciurus hudsonicus*) tend to have consistent male biased ASR (Hurly 1987). Variation in ASR may be due to four non-mutually exclusive factors (Wilson 1975; Székely et al. 2014a): first, biased ASR may emerge as a consequence of biased juvenile sex ratios. Sex ratios may already be biased at conception (primary sex ratio), or at birth or hatching (secondary sex ratio), or male and female juveniles may die at different rates (Wilson 1975; Clutton-Brock 1991; Kalmbach and Benito 2007). Second, differential adult survival may bias ASR, because one sex may be predated more often than the other (Berger and Gompper 1999; Hirst et al. 2010), or be more susceptible to parasites and pathogens (Moore and Wilson 2002; Valdebenito et al. 2021). Third, maturation often depends on growth rates, and if one sex is larger than the other, the larger sex may take longer to mature, as is the case in raptors or mammals (Promislow 1992; Benito and Gonzáles-Solís 2007; Hirst et al. 2010). Fourth, emigration and immigration are often sex-specific. For instance in birds, males tend to stay in their natal territory and females disperse, whereas in mammals the reverse tends to be the case (Clobert et al. 2012; Végvári et al. 2018). While ASR variation seems to be influenced by demographic factors, still there are important information gaps in terms of ecological and life history variables, particularly at specific taxonomic levels.

Shorebirds and allies (Scolopaci and Charadrii, sandpipers, plovers and allies) are particularly interesting from a sexual selection perspective since they show an unusually great diversity in mating strategies and sexual differences in behaviour, size and colouration (Székely 2019). For instance, despite being a relatively small group (currently, about 219 species; Gill et al. 2023), their members display examples for extreme polygyny like the Ruff (*Calidris pugnax*), while others show reversed size dimorphism and sex role reversal like most jacanas (Figuerola 1999; Fresneau et al. 2021). Shorebirds also typically nest on the ground in open-cup nests, are precocial, and often show distinctive long distance movements, which can contribute for additional selection pressures of e.g. colouration, size and shape (Hedenström 2008; Stevens and Merilaita 2009). Interestingly, shorebirds show substantial ASR variation and recent species-specific studies have suggested that sex-specific mortality occurring in life stages prior to adulthood is an important contributor to ASR biases (Eberhart-Phillips et al. 2017; Eberhart-Phillips et al. 2018; Loonstra et al. 2019). It seems that a female-biased mortality during the chick stage would likely translate into a male-biased ASR, and vice versa (Eberhart-Phillips et al. 2018). This supports theoretical and empirical work that identify sex-specific mortality as a driver of ASR variation (Stenzel et al. 2011; Székely et al. 2014b). However, it is unknown which variables determine sex-specific mortality in shorebirds. On the other hand, sex roles, that describe sex differences in courtship, mate competition, social pair-bonds and parental care, seem to be heavily influenced by ASR variation in birds (Liker et al. 2013; Liker et al. 2021). Although Liker et al. (2013) showed that parental care and mating system were associated with ASR in shorebirds, this relationship is still yet to be explored in other sex role components that are usually not well described in shorebirds (i.e. there is substantial missing data) and thus their degree of sex bias is unknown.

In this work we used phylogenetic comparative methods to examine the predictors of ASR variation across shorebirds. This was achieved by collecting data from the literature from birds of the suborder Charadrii and Scolopaci (n = 205 species) on ASR, sex role components and sex-specific adult mortality. Additionally, we also collected data on hatching sex ratio (Benito and Gonzáles-Solís 2007). Among the hypotheses that connect the aforementioned variables, we note the energetic costs incurred by the parents when producing the larger offspring in sexually dimorphic species; sex-specific mortality at different age stages being a driver of ASR; and abundance of the common sex serving as a likely resource for care of the offspring (Fisher 1934; Székely et al. 2014b; Gonzalez-Voyer et al. 2022). First, we determined the overall sex-specific distribution of each variable to determine their degree of sex bias. We then conducted phylogenetically informed model analysis to evaluate how ASR and sex-specific mortality was associated to each variable, including sexual size dimorphism, sexual dichromatism, mating system and parental care. Because hatching sex ratio can be important for determining sex ratios later in life (Eberhart-Phillips et al. 2018; Schacht et al. 2022), we also used a subset of the data to explore its relevance in some of the aforementioned associations.

## Methods

### Data collection

We used the reference list provided by the IOC World Bird List (Gill et al. 2023) to obtain the most up-to-date list of species from the suborders Scolopaci and Charadrii, hereafter referred to as shorebirds. This list had 219 birds, including six extinct species, one rare, critically endangered species with very little information available, and other seven species that in recent years had been considered as subspecies. These 14 taxa were removed from further analysis (Supplemental Material Table S1), either because they were not included in the latest phylogenetic hypothesis, and/or because there was no available information on their appearance and biology –apart from hand-made sketches or pictures of museums specimens. Thus, the final list of species used in the analysis had 205 shorebird species from 13 families (Supplemental Material Figure S1).

For data collection we used as starting point the dataset provided by Székely et al. (2022), containing sexual selection and life-history data for 9,982 wild bird species, nearly all living birds on the planet. These data spanned until 2015, therefore to obtain the most up-to-date and comprehensive dataset we updated it up to February 2023 by conducting a systematic literature search following the PRISMA scheme (Moher et al. 2009), using ISI Web of Science (see chart in Supplementary Material Figure S2). We also consulted publicly available sources including books, previous data compilations (such as Méndez et al. 2018; and Storchová and Hořák 2018), the Birds of the World (The Cornell Ornithology Lab; Billerman et al. 2022), and experts on particular species. Our inclusion criteria required these data to be: (i) determined from birds of known sex (molecular or morphological sexing), (ii) obtained from free-living wild birds (not captive), (iii) from populations that were not experimentally manipulated, and (iv) both males and females should have belonged to the same population and sampled during the same timeframe.

### Estimating variables

*ASR and hatching sex ratio*: ASR was calculated as the proportion of adult males in the adult population (Liker et al. 2013; Székely et al. 2014a). Hatching sex ratio corresponded to the proportion of male chicks in a brood population. Estimates were based on a variety of methods (e.g. censuses of individually marked breeding adults, demographic modelling, counts of captured and dead birds). ASR estimates are repeatable between populations of the same species as measured by the intraclass correlation coefficient (ICC = 0.64; Ancona et al. 2017). Moreover, by including several ASR estimates from different population of the same species, we overcome other possible sources of variation such as latitude (Nebel et al. 2002). Referring to hatching sex ratio, Fiala (1980) warned that excluding broods that had suffered losses prior to sexing had the potential to bias the sample in favour of the sex with greater survivorship. If broods that had suffered losses (incomplete broods) did not exhibit a hatching sex ratio differing significantly from complete broods, we followed the convention of Fiala (1981) and included sex ratio data of both complete and incomplete broods.

*Sex-specific mortality:* this variable considered estimates of males and females only for the age classes after the age of first reproduction. Data corresponded to either sex-specific annual mortality or annual survival (φ). In the latter case, we calculated mortality by simply subtracting the survival estimate to 1 (i.e. 1-φ). Adult annual survival may be estimated by various methods (return rates, mark-recapture or demographic analyses), although we used better quality estimates (e.g. those from mark-recapture analyses) whenever we had a choice. Sex-specific mortality was calculated as male annual mortality minus female annual mortality.

*Sex bias in parental care:* we collected data of both incubation and brooding, two of the main parental care behaviours in shorebirds since chick feeding is not globally present in this avian group. Both variables were scored separately using a 5-point score of the relative effort of the male, where: 0 = no involvement of male; 1 = 1-33%; 2 = 34-66%; 3 = 67-99%; 4 = 100%. Parental care bias was considered as the mean score between incubation and brooding. This variable was later scaled in order to set equal parental care contribution of the sexes to zero.

*Sexual size dimorphism (SSD):* this variable was computed as log(adult male body mass (g)/adult female body mass (g)).

*Sexual dichromatism:* we used the plumage scoring system provided by Gonzalez-Voyer et al. (2022; full details therein). In short, the bird was split into 5 body regions (head; nape, back and rump; throat, chest and belly; tail; and wings). For a given species, each body region was scored separately with a score system that ranged between –2 and 2, according to the following: –2, the female was substantially brighter and/or more patterned than the male; –1, the female was brighter and/or more patterned than the male; 0, there was no difference in the body region or there was difference but neither could be considered brighter than the other; 1, the male was brighter and/or more patterned than the female; 2, the male was substantially brighter and/or more patterned than the female. For the analysis we used the mean of these five regions.

*Polygamy score:* we used the difference between scores of female polygamy and male polygamy as a proxy for the social mating system based on the scoring system in Liker et al. (2015). The overall incidence of polygamy for each sex were expressed on a 5-point score, with 0 corresponding to no (or very rare) polygamy (< 0.1% of individuals), 1 to rare polygamy (0.1–1%), 2 to uncommon polygamy (1–5%), 3 to moderate polygamy (5–20%), and 4 to common polygamy (> 20%; including males in lekking species to express the high variance in male mating success in these species; Höglund and Alatalo 1995). We calculated the sex bias in polygamy score as male score minus female score.

### Statistical analysis

The structure of the analysis followed three steps involving phylogenetic comparative analyses. First, we determined the overall distribution of each variable, that is how much it deviated from zero (i.e. no sex difference). Second, we studied the level of association among sex role components. Lastly, we investigated which variables predicted ASR and sex-specific mortality. To learn about the distribution of each variable as well as investigating whether they deviated from parity, we built generalised linear mixed models, with each variable as response variable and run to the intercept. For this purpose, we used the R package MCMCglmm (Hadfield 2010). This tests whether the mean estimate of a given variable is significantly different from zero. To evaluate hypothesised correlations among sex role components (Janicke et al. 2016; Mokos et al. 2021; Gonzalez-Voyer et al. 2022), we conducted marginal correlation accounting for phylogenetic relatedness using the *mvBM()* function, from the R package mvMORPH (Clavel et al. 2015).

Regression analysis were conducted by building models with ASR or sex-specific mortality as response variable, and SSD, sexual dichromatism, sex bias in polygamy and the sex bias in parental care as explanatory variables. Hatching sex ratio was an explanatory variable only for the ASR analysis. One last analysis was conducted having ASR as response variable and sex-specific mortality as explanatory variable. Phylogeny (a variance– covariance matrix) and species (to correct for species for which we had several estimates) were added as random-effect variables. We used the avian phylogeny proposed by (Jetz et al. 2012), using a consensus tree from 1,000 randomly selected trees from a pool of 10,000 available trees (http://birdtree.org). The consensus tree was computed with the function *ls*.*consensus()* from the R package phytools, which computes the least-squares consensus tree from the mean patristic distance matrix of a set of trees (Revell 2012). We used parameter expanded (random-effects) and inverse-Wishart priors (fixed-effects) based on improving model convergence. Family distribution in all models was Gaussian. The number of iterations, thinning and burn-in was set at 501,000, 500, and 1,000, respectively, in order to obtain an effective sample size of 1,000. Convergence and autocorrelation levels were assessed through the Gelman-Rubin test (Gelman and Rubin 1992), trace graphs and the *autocorr()* function, implemented in the R package CODA (Plummer et al. 2006). MCMCglmm results are expressed as posterior mean, lower and upper 95% credible intervals, and significance as a pMCMC value. All statistical analyses were conducted in the R statistical software version 4.2.2 (R Core Team 2023).

## Results

### Overall distributions of variables

We found strong sex biases across the seven variables investigated, however, none of them significantly deviated from parity (i.e. different from zero, Figure 1). The two most sex-biased variables were ASR and sex-specific mortality with a male and a female bias, respectively (Figure 1).

**FIGURE 1.**
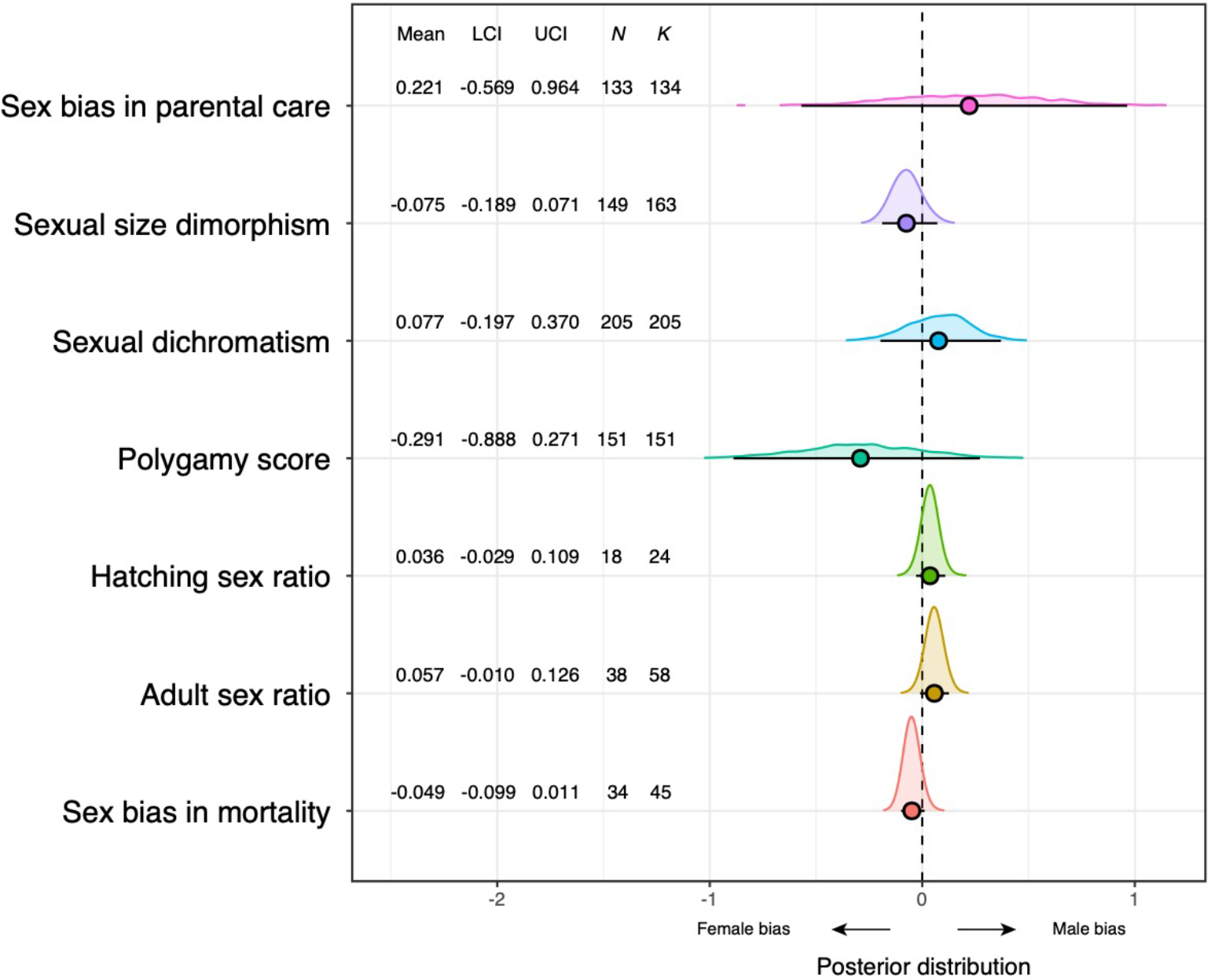
Sex-specific distribution of seven variables in shorebirds. The distributions were built by plotting the density of the posterior distribution of MCMCglmm models run to the intercept. Each variable has a mean (coloured circle), 95% lower and upper credible intervals (LCI, UCI) and their density on top. *N* refers to the number of species and *K* to the number of individual estimates for a given variables (i.e. some species had more than one estimate).

### Correlation among sex role components

Overall, the four sex role components showed rather strong correlations. Sexual dichromatism was the variable having the weakest correlation coefficients, being only strongly correlated with SSD. All other variables had moderate to strong correlation coefficients (Figure 2a). Noticeably, parental care negatively correlates with the other variables (Figure 2a), suggesting that when one sex is brighter, larger or more polygamous, the other sex will provide most of the parental care (e.g. Jacanas group, see Figure 2b).

**FIGURE 2.**
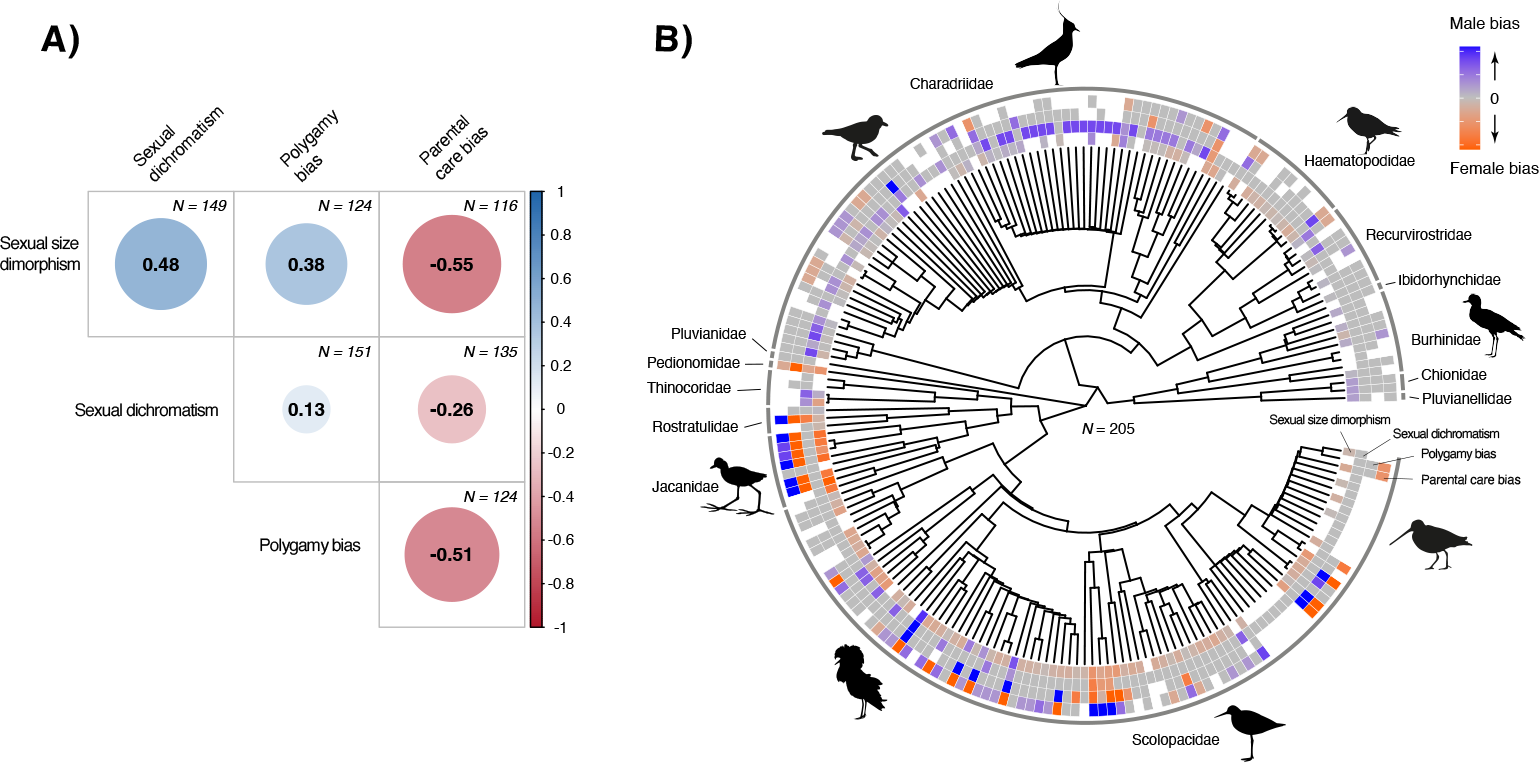
(**A**) Phylogenetically controlled correlation matrix of four sex role components in shorebirds. Numbers in centre of circles indicate the value of the phylogenetically controlled correlation, also depicted by the size and colour of the circles. *N* indicates the number of species in each specific bivariate correlation. (**B**) Distribution of sex role components in the shorebird tree (*N* = 205). Heatmap represents degree of sex bias, where the colour grey (i.e. a value of zero) denotes no sex bias. White squares represent lack of data for a given variable and species.

### Predictors of ASR and sex-specific mortality

ASR was significantly predicted by all sex role components, except for sexual dichromatism (Table 1). The direction of the association was negative in all models except for parental care bias, where a positive slope indicates that increases in male-biased ASR was associated with increased male care, and vice versa (Figure 3). Sex-specific mortality was not significantly predicted by any variable (Table 1).

**TABLE 1.** Adult sex ratio and sex bias in mortality in relation to different sex-specific predictors. Mean refers to the mean posterior distribution of MCMCglmm models. LCI and UCL are lower and upper 95% credible intervals. *K* is the number of individual estimates for a given variables, which in this case can also be interpreted as sample size. Bold font indicates statistical significance.

| | $K$ | Mean | LCI | UCL | pMCMC |
| --- | --- | --- | --- | --- | --- |
| Adult sex ratio |  |  |  |  |  |
| **Sexual size dimorphism** | **64** | **-0.250** | **-0.451** | **-0.071** | **0.018** |
| Sexual dichromatism | 59 | -0.033 | -0.119 | 0.049 | 0.434 |
| **Sex bias in polygamy** | **56** | **-0.038** | **-0.069** | **-0.007** | **0.024** |
| **Sex bias in parental care** | **55** | **0.037** | **0.004** | **0.069** | **0.034** |
| Hatching sex ratio | 21 | -1.007 | -2.275 | 0.359 | 0.122 |
| Sex bias in mortality | 35 | -0.014 | -0.495 | 0.441 | 0.952 |
| Sex bias in mortality |  |  |  |  |  |
| Sexual size dimorphism | 50 | -0.113 | -0.330 | 0.121 | 0.336 |
| Sexual dichromatism | 47 | -0.061 | -0.150 | 0.027 | 0.164 |
| Sex bias in polygamy | 46 | 0.023 | -0.017 | 0.068 | 0.298 |
| Sex bias in parental care | 46 | -0.030 | -0.084 | 0.020 | 0.266 |

**FIGURE 3.**
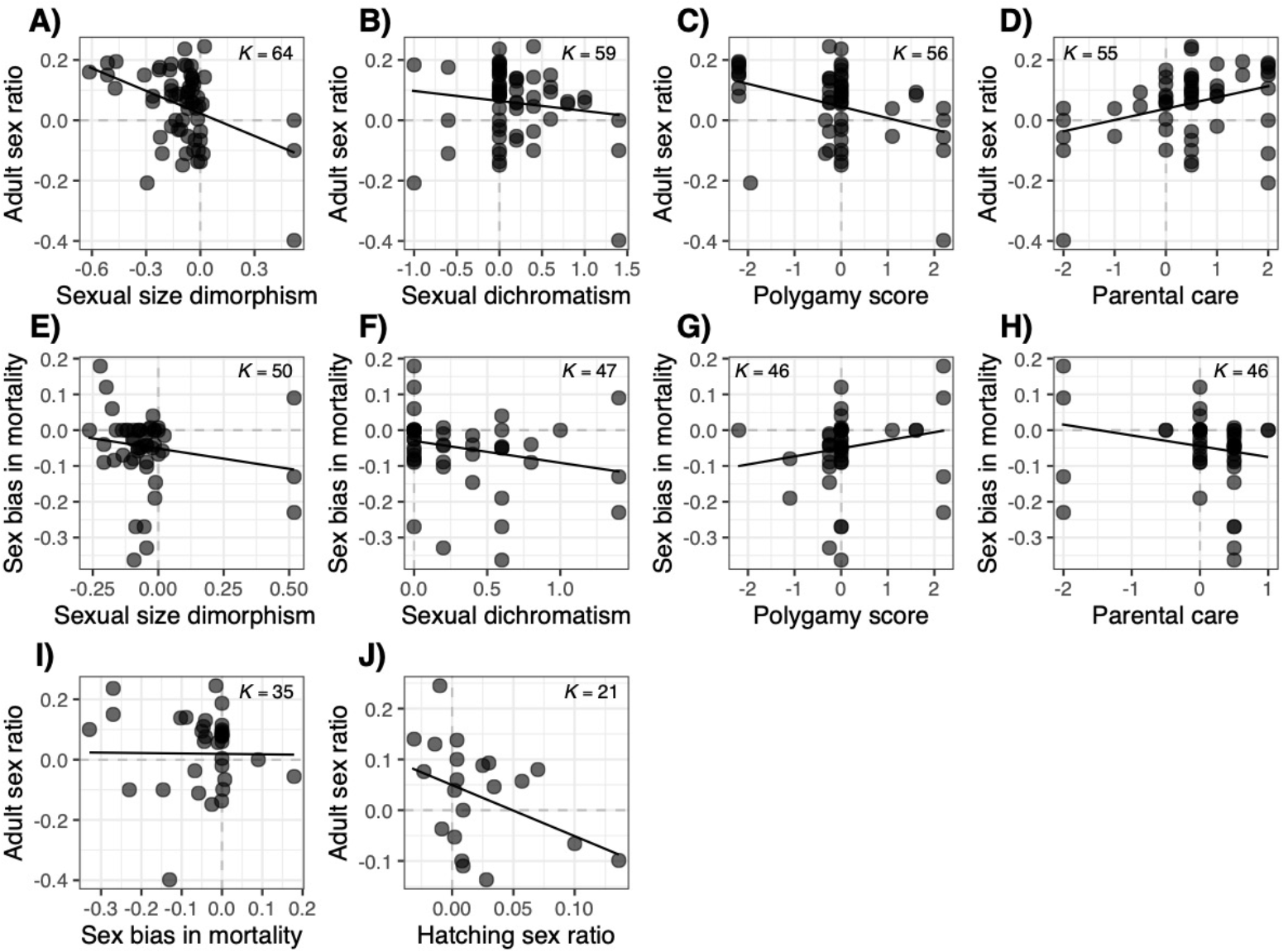
Adult sex ratio and sex bias in mortality in relation to sex role components in shorebirds (**A-D** and **E-H**, respectively). Adult sex ratio is also tested against hatching sex ratio and the sex bias in mortality (**I** and **J**). Datapoints correspond to raw data and regression lines were extracted from MCMC generalised linear models. In all plots, zero means no sex difference (dashed line), positive values represent a male bias and negative values a female bias. *K* refers to the number of individual estimates for a given analysis, which in this case can also be interpreted as sample size.

## Discussion

In this work we compiled extensive information on life-history, sex roles and demographic variables of male and female shorebirds, in order to, to our knowledge, provide the most comprehensive characterisation of ASR and sex-specific predictors in a bird group with high diversity in mating strategies and sexual differences. Our analysis showed important sex-specific variation in most variables studied, where we highlight a trend towards sex role reversal. ASR was broadly predicted by sex role components but, interestingly, these variables were not associated with sex-specific mortality.

Deviation from sex-specific parity in sexual selection, demographic and sex role variables are not uncommon in nature. For instance, Donald (2007) found a significant male bias in ASR across wild birds, whereas Lemaître et al. (2020) found that lifespan in mammals was female-biased. Our results showed no significant sex bias in the seven variables here studied though we noted strong sex-specific trends. Shorebird sex ratios tended to be already male-biased at hatching, which seems to be maintained when they reach adulthood. If we also take into consideration that in most species females were larger than males, this is consistent with Fisher–Trivers’ frequency-dependent argument that in sexually dimorphic species parents should produce more of the ‘cheaper’ sex (i.e. males) (Trivers 1985). In particular, we found that, overall, parental care, polygamy score and sexual size dimorphism did not follow the conventional picture known from sex role theory but rather a sex role reversal (Janicke et al. 2016). These four sex role components also correlated in a direction opposite to the expected and previously shown in birds (Gonzalez-Voyer et al. 2022). It has been well acknowledged that ASR variation is related to the evolution of sex roles, including sex role reversal (Liker et al. 2013), however it is still unknown the likely evolutionary path that drove the evolution of each trait. One interpretation of sexual selection theory predicts that males obtain greater fitness benefits than females through multiple mating because sperm is cheaper to produce than eggs, thus sexual selection should lead to the evolution of male-biased secondary sexual characters (Bateman 1948). We saw that in shorebirds, colouration and size dimorphism showed strong covariation, but females tend to be larger than males. These findings contrast ideas drawn from sexual selection theory but support a recent study by Mokos et al. (2021) that found that anisogamy was not globally associated with the expected direction of sexual dimorphism. Our results also contrast the findings of Gonzalez-Voyer et al. (2022) across the avian tree of life, where they found no correlation between SSD and sexual dichromatism (correlation coefficient = 0.05). In our work most sex role components moderately or strongly correlated with each other, suggesting strong sexual selection forces acting upon these traits in shorebirds. The negative correlation between the sex bias in parental care and the other sex role variables shows that parental care is mostly provided by the duller, smaller, and/or less polygamous sex, regardless of that sex being a male or a female.

ASR was predicted by SSD, parental care bias and polygamy bias, supporting previous work by Székely et al. (2014b) and Liker et al. (2021). ASR was not associated with hatching sex ratio, in accordance with Székely et al. (2014b). While some species with a male-biased offspring sex ratio may exhibit male-biased ASR (e.g. Kentish plover *C. alexandrinus*; Kosztolányi et al. 2011), and other species with female-biased offspring sex ratio may exhibit female-biased ASR (Ruff *Calidris pugnax*; Jaatinen et al. 2010), the lack of a significant relationship suggests that the pattern is not globally present across shorebirds. Similar conclusions can be inferred from the lack of association between ASR and sex-specific mortality, contrary to the findings of Székely et al. (2014b). Studies addressing sex-specific mortality from a comparative perspective are rather scarce. The work of Promislow (1992), Liker and Székely (2005), Székely et al. (2014b) and Méndez et al. (2018) showed that sex-specific mortality in birds was strongly correlated with, for example, wing size dimorphism, ASR, SSD, mating competition and post-hatching parental care. In our work we found no support for these previously found associations. Perhaps the particular phenology and ecology of shorebirds enables mortality events to occur by means independent from their sex roles. For example, in migratory species –as many shorebirds–, the competing sex tends to arrive earlier than the other in order to define their territories in anticipation of the arrival of the other sex (such as protandry; Kokko et al. 2006). This sex difference in departure from non-breeding grounds and/or arrival to breeding grounds, as well as a potentially different propensity to abort migration one year to improve survival (Tavera et al. 2020), could drive additional sex differences in mortality. Therefore, it is possible that in this bird group different sexual mortality patterns cannot be explained solely by their sex roles (Palacín et al. 2009; Bosman et al. 2012; Macdonald et al. 2016).

Shorebirds are an intriguing avian group that seems to have evolved sex roles that are not often the norm for most avian groups (Gonzalez-Voyer et al. 2022). Previous studies have linked sex role reversal with ASR variation but further discussion as to the evolutionary origins of such qualities are still contested, arguing that much care should be taken when attempting to interpret the work of Bateman (1948), particularly of anisogamy as driver of conventional sex roles (Janicke et al. 2016). Nevertheless, sex roles components are consistent predictors of ASR in birds, including shorebirds. This suggests that their implications in shorebird’s life history are substantial and therefore distorters of a species’ sex ratio could potentially entail a cascade of events affecting their behaviour, ecology and, ultimately, their conservation. An example occurs in Kentish plover, where ASRs may contrast substantially between populations, which are usually accompanied by their corresponding contrasting sex-skew in parental care behaviour (Amat et al. 1999; Carmona-Isunza et al. 2017; Que et al. 2019).

Despite female shorebirds tending to die at higher rates than males, sex-specific mortality had no association with the variables here tested. We note the complex life-cycles of many shorebirds which could involve variables other than life history that we have not considered. Nonetheless, the relatively good coverage of our dataset enabled us to unveil intriguing relationships not often seen in birds. Our results confirm that sex role components are importantly associated with ASR variation and suggest that in shorebirds these traits are under stronger sexual selection pressures. Finally, we take value on conducting highly detailed studies on well-known species. However, we encourage researchers to expand their coverage to species with little knowledge available, such as Neotropical shorebirds. Hopefully in a few years, a future phylogenetic tree displaying shorebird sex roles will have no missing data and, therefore, a refined analysis will shed additional light on the drivers of ASR variation in shorebirds.

## Acknowledgements

We thank Narhulan Halimbekh for providing sex-specific body mass data on the Lesser sand plover (*Charadrius mongolus*), and Nolwenn Fresneau for providing ASR data from Pheasant-tailed jacana (*Hydrophasianus chirurgus*). Funding statement: JOV was funded by ANID Fondecyt postdoc ID: 3220722. JGN was funded by ANID Millennium Science Initiative Program—ICN2021_002. Conflict of interest statement: the authors declare no competing interests. Author contributions: JOV conceived the idea and design of study, as well as conducted the data analysis. TT collected the data. JGN supervised the research. All authors contributed substantially to revisions of the paper.

## Literature cited

Amat, J. A., R. M. Fraga, and G. M. Arroyo (1999). Brood desertion and polygamous breeding in the Kentish Plover Charadrius alexandrinus. Ibis 141:596–607.

Ancona, S., F. V. Dénes, O. Krüger, T. Székely, and S. R. Beissinger (2017). Estimating adult sex ratios in nature. Philosophical Transactions of the Royal Society B 372: 20160313.

Bateman, A. J. (1948). Intra-sexual selection in Drosophila. Heredity 2:349–368.

Benito, M. M., and J. Gonzáles-Solís (2007). Sex ratio, sex-specific chick mortality and sexual size dimorphism in birds. Journal of Evolutionary Biology 20:1522–1530.

Berger, J., and M. E. Gompper (1999). Sex Ratios in Extant Ungulates: Products of Contemporary Predation or Past Life Histories? Journal of Mammalogy 80:1084–1113.

Billerman, S. M., B. K. Keeney, P. G. Rodewald, and T. S. Schulenberg (2022). Birds of the World. Available at: https://birdsoftheworld.org/bow/home.

Bosman, D. S., H. J. P. Vercruijsse, E. W. M. Stienen, M. Vincx, L. De Neve, and L. Lens (2012). Effects of body size on sex-related migration vary between two closely related gull species with similar size dimorphism. Ibis 154:52–60.

Carmona-Isunza, M. C., S. Ancona, T. Székely, A. P. Ramallo-González, M. Cruz-López, M. A. Serrano-Meneses, and C. Küpper (2017). Adult sex ratio and operational sex ratio exhibit different temporal dynamics in the wild. Behavioral Ecology 28:523–532.

Clavel, J., G. Escarguel, and G. Merceron (2015). mvmorph: an r package for fitting multivariate evolutionary models to morphometric data. Methods in Ecology and Evolution 6:1311–1319.

Clobert, J., M. Baguette, T. G. Benton, and J. M. Bullock. 2012. Dispersal ecology and evolution. Oxford University Press, Oxford, UK.

Clutton-Brock, T. H. 1991. The evolution of parental care. Princeton University Press, Princeton, NJ.

Donald, P. F. (2007). Adult sex ratios in wild bird populations. Ibis 149:671–692.

Eberhart-Phillips, L. J., C. Küpper, T. E. X. Miller, M. Cruz-López, K. H. Maher, N. dos Remedios, M. A. Stoffel, J. I. Hoffman, O. Krüger, and T. Székely (2017). Sex-specific early survival drives adult sex ratio bias in snowy plovers and impacts mating system and population growth. Proceedings of the National Academy of Sciences 114:E5474–E5481.

Eberhart-Phillips, L. J., C. Küpper, M. C. Carmona-Isunza, O. Vincze, S. Zefania, M. Cruz-López, A. Kosztolányi, T. E. X. Miller, Z. Barta, I. C. Cuthill, T. Burke, et al. (2018). Demographic causes of adult sex ratio variation and their consequences for parental cooperation. Nature Communications 9:1651.

Fiala, K. L. (1980). On Estimating the Primary Sex Ratio from Incomplete Data. The American Naturalist 115:442–444.

Fiala, K. L. (1981). Sex Ratio Constancy in the Red-Winged Blackbird. Evolution 35:898–910.

Figuerola, J. (1999). A comparative study on the evolution of reversed size dimorphism in monogamous waders. Biological Journal of the Linnean Society 67:1–18.

Fisher, R. 1934. The genetic theory of natural selection. Oxford University Press, Oxford, UK.

Fresneau, N., Y.-F. Lee, W.-C. Lee, A. Kosztolányi, T. Székely, and A. Liker (2021). Sex Role Reversal and High Frequency of Social Polyandry in the Pheasant-Tailed Jacana (Hydrophasianus chirurgus). Frontiers in Ecology and Evolution 9:742588.

Gelman, A., and D. B. Rubin (1992). Inference from Iterative Simulation Using Multiple Sequences. Statistical Science 7:457–472.

Gill, F., D. Donsker, and P. Rasmussen. 2023. IOC World Bird List (v13.1).

Gonzalez-Voyer, A., G. H. Thomas, A. Liker, O. Krüger, J. Komdeur, and T. Székely (2022). Sex roles in birds: Phylogenetic analyses of the influence of climate, life histories and social environment. Ecology Letters 25:647–660.

Gusmão, L. F. M., and A. D. McKinnon (2009). Sex ratios, intersexuality and sex change in copepods. Journal of Plankton Research 31:1101–1117.

Hadfield, J. D. (2010). MCMC Methods for Multi-Response Generalized Linear Mixed Models: The MCMCglmm R Package. Journal of Statistical Software 33:1–22.

Hays, G. C., A. D. Mazaris, G. Schofield, and J.-O. Laloë (2017). Population viability at extreme sex-ratio skews produced by temperature-dependent sex determination. Proceedings of the Royal Society B: Biological Sciences 284:20162576.

Hedenström, A. (2008). Adaptations to migration in birds: behavioural strategies, morphology and scaling effects. Philosophical Transactions of the Royal Society B: Biological Sciences 363:287–299.

Hirst, A. G., D. Bonnet, D. V. P. Conway, and T. Kiørboe (2010). Does predation controls adult sex ratios and longevities in marine pelagic copepods? Limnology and Oceanography 55:2193–2206.

Höglund, J., and R. V. Alatalo. 1995. Leks. Princeton University Press, New Jersey.

Hurly, T. A. (1987). Male-biased adult sex ratios in a red squirrel population. Canadian Journal of Zoology 65:1284–1286.

Jaatinen, K., A. Lehikoinen, and D. B. Lank (2010). Female-biased sex ratios and the proportion of cryptic male morphs of migrant juvenile Ruffs (Philomachus pugnax) in Finland. Ornis Fennica 87:125–134.

Janicke, T., I. K. Häderer, M. J. Lajeunesse, and N. Anthes (2016). Darwinian sex roles confirmed across the animal kingdom. Science Advances 2:e1500983.

Jetz, W., G. H. Thomas, J. B. Joy, K. Hartmann, and A. O. Mooers (2012). The global diversity of birds in space and time. Nature 491:444–448.

Kalmbach, E., and M. M. Benito (2007). Sexual size dimorphism and offspring vulnerability in birds. In Sex, size and gender roles (F. D.J., B. W.U., and S. T., Editors). Oxford University Press, Oxford, UK:133–142.

Kokko, H., T. G. Gunnarsson, L. J. Morrell, and J. A. Gill (2006). Why do female migratory birds arrive later than males? Journal of Animal Ecology 75:1293–1303.

Kosztolányi, A., Z. Barta, C. Küpper, and T. Székely (2011). Persistence of an extreme male-biased adult sex ratio in a natural population of polyandrous bird. Journal of Evolutionary Biology 24:1842–1846.

Lemaître, J.-F., V. Ronget, M. Tidière, D. Allainé, V. Berger, A. Cohas, F. Colchero, D. A. Conde, M. Garratt, A. Liker, G. A. B. Marais, et al. (2020). Sex differences in adult lifespan and aging rates of mortality across wild mammals. Proceedings of the National Academy of Sciences 117:8546–8553.

Liker, A., and T. Székely (2005). Mortality costs of sexual selection and parental care in natural populations of birds. Evolution 59:890–897.

Liker, A., R. P. Freckleton, and T. Székely (2013). The evolution of sex roles in birds is related to adult sex ratio. Nature Communications 4:1587.

Liker, A., R. P. Freckleton, V. Remeš, and T. Székely (2015). Sex differences in parental care: Gametic investment, sexual selection, and social environment. Evolution 69:2862–2875.

Liker, A., V. Bókony, I. Pipoly, J. F. Lemaître, J. M. Gaillard, T. Székely, and R. P. Freckleton (2021). Evolution of large males is associated with female-skewed adult sex ratios in amniotes. Evolution 75:1636–1649.

Loonstra, A. H. J., M. A. Verhoeven, N. R. Senner, J. C. E. W. Hooijmeijer, T. Piersma, and R. Kentie (2019). Natal habitat and sex-specific survival rates result in a male-biased adult sex ratio. Behavioral Ecology 30:843–851.

Macdonald, C. A., E. A. McKinnon, H. G. Gilchrist, and O. P. Love (2016). Cold tolerance, and not earlier arrival on breeding grounds, explains why males winter further north in an Arctic-breeding songbird. Journal of Avian Biology 47:7–15.

Méndez, V., J. A. Alves, J. A. Gill, and T. G. Gunnarsson (2018). Patterns and processes in shorebird survival rates: a global review. Ibis 160:723–741.

Moher, D., A. Liberati, J. Tetzlaff, and D. G. Altman (2009). Preferred reporting items for systematic reviews and meta-analyses: the PRISMA statement. PLoS Med 6:e1000097.

Mokos, J., I. Scheuring, A. Liker, R. P. Freckleton, and T. Székely (2021). Degree of anisogamy is unrelated to the intensity of sexual selection. Scientific Reports 11:19424.

Moore, S. L., and K. Wilson (2002). Parasites as a Viability Cost of Sexual Selection in Natural Populations of Mammals. Science 297:2015–2018.

Nebel, S., D. B. Lank, P. D. O’Hara, G. Fernández, B. Haase, F. Delgado, F. A. Estela, L. J. E. Ogden, B. Harrington, B. E. Kus, J. E. Lyons, et al. (2002). Western Sandpipers (Calidris Mauri) During the Nonbreeding Season: Spatial Segregation on a Hemispheric Scale. The Auk 119:922–928.

Palacín, C., J. C. Alonso, J. A. Alonso, C. A. Martín, M. Magaña, and B. Martin (2009). Differential Migration by Sex in the Great Bustard: Possible Consequences of an Extreme Sexual Size Dimorphism. Ethology 115:617–626.

Plummer, M., N. Best, K. Cowles, and K. Vines (2006). CODA: convergence diagnosis and output analysis for MCMC. R News 6:7–11.

Promislow, D. E. L. (1992). Costs of sexual selection in natural populations of mammals. Proceedings of the Royal Society of London. Series B: Biological Sciences 247:203–210.

Que, P., T. Székely, P. Wang, Q. Lu, W. Lei, Y. Liu, and Z. Zhang (2019). Offspring sex ratio is unrelated to parental quality and time of breeding in a multiple-breeding shorebird. Journal of Ornithology 160:443–452.

Revell, L. J. (2012). phytools: an R package for phylogenetic comparative biology (and other things). Methods in Ecology and Evolution 3:217–223.

Schacht, R., S. R. Beissinger, C. Wedekind, M. D. Jennions, B. Geffroy, A. Liker, P. M. Kappeler, F. J. Weissing, K. L. Kramer, T. Hesketh, J. Boissier, et al. (2022). Adult sex ratios: causes of variation and implications for animal and human societies. Communications Biology 5:1273.

Stenzel, L. E., G. W. Page, J. C. Warriner, J. S. Warriner, K. K. Neuman, D. E. George, C. R. Eyster, and F. C. Bidstrup (2011). Male-skewed adult sex ratio, survival, mating opportunity and annual productivity in the Snowy Plover Charadrius alexandrinus. Ibis 153:312–322.

Stevens, M., and S. Merilaita (2009). Animal camouflage: current issues and new perspectives. Philosophical Transactions of the Royal Society B: Biological Sciences 364:423–427.

Storchová, L., and D. Hořák (2018). Life-history characteristics of European birds. Global Ecology and Biogeography 27:400–406.

Székely, T. (2019). Why study plovers? The significance of non-model organisms in avian ecology, behaviour and evolution. Journal of Ornithology 160:923–933.

Székely, T., F. J. Weissing, and J. Komdeur (2014a). Adult sex ratio variation: implications for breeding system evolution. Journal of Evolutionary Biology 27:1500–1512.

Székely, T., A. Liker, R. P. Freckleton, C. Fichtel, and P. M. Kappeler (2014b). Sex-biased survival predicts adult sex ratio variation in wild birds. Proceedings of the Royal Society B: Biological Sciences 281:20140342.

Székely, T., A. Liker, G. H. Thomas, J. Komdeur, O. Krüger, and A. Gonzalez-Voyer (2022) Sex roles in birds: influence of climate, life histories and social environment, Dryad, Dataset, 10.5061/dryad.fbg79cnw7

Tavera, E. A., G. E. Stauffer, D. B. Lank, and R. C. Ydenberg (2020). Oversummering juvenile and adult Semipalmated sandpipers in Perú gain enough survival to compensate for foregone breeding opportunity. Movement Ecology 8:42.

R Core Team (2023). R: A language and environment for statistical computing. R Foundation for Statistical Computing Vienna, Austria.

Trivers, R. 1985. Social evolution. The Benjamin/Cummings Publishing Company, Menlo Park, CA.

Valdebenito, J. O., N. Halimubieke, Á. Z. Lendvai, J. Figuerola, G. Eichhorn, and T. Székely (2021). Seasonal variation in sex-specific immunity in wild birds. Scientific Reports 11:1349.

Végvári, Z., G. Katona, B. Vági, R. P. Freckleton, J.-M. Gaillard, T. Székely, and A. Liker (2018). Sex-biased breeding dispersal is predicted by social environment in birds. Ecology and Evolution 8:6483–6491.

Wilson, E. O. 1975. Sociobiology: the new synthesis. Harvard University Press, Cambridge, MA.

