## Supplemental file for "Adult sex ratio variation and its sex-specific predictors in shorebirds"

**Table S1**. Summary of inclusion criteria of 219 shorebird species according to their information available.

|  | Species | Conservation status | Notes |
| --- | --- | --- | --- |
| Extinct species | | | |
| 1 | *Coenocorypha barrierensis* | EX |  |
| 2 | *Coenocorypha iredalei* | EX |  |
| 3 | *Haematopus meadewaldoi* | EX |  |
| 4 | *Prosobonia cancellate* | EX |  |
| 5 | *Prosobonia ellisi* | EX |  |
| 6 | *Prosobonia leucoptera* | EX |  |
| Critically endangered species with no data | | | |
|  | *Vanellus macropterus* | CR | no info |
| Species considered as subspecies of species already included | | | |
| 1 | *Burhinus indicus* | LC | considered subspecies of *Burhinus oedicnemus* |
| 2 | *Charadrius dealbatus* | LC | considered subspecies of *Charadrius alexandrinus* |
| 3 | *Coenocorypha huegeli* | NT | considered subspecies of *Coenocorypha aucklandica* |
| 4 | *Gallinago delicata* | LC | considered subspecies of *Gallinago gallinago* |
| 5 | *Gallinago magellanica* | LC | considered subspecies of *Gallinago paraguaiae* |
| 6 | *Himantopus melanurus* | LC | considered subspecies of *Himantopus mexicanus* |
| 7 | *Numenius hudsonicus* | LC | considered subspecies of *Numenius phaeopus* |
| Corrected species | |  |  |
| 1 | *Prosobonia parvirostris* | EN | replaced for *Prosobonia cancellata* |
| Total species in analysis = 205 | | | |

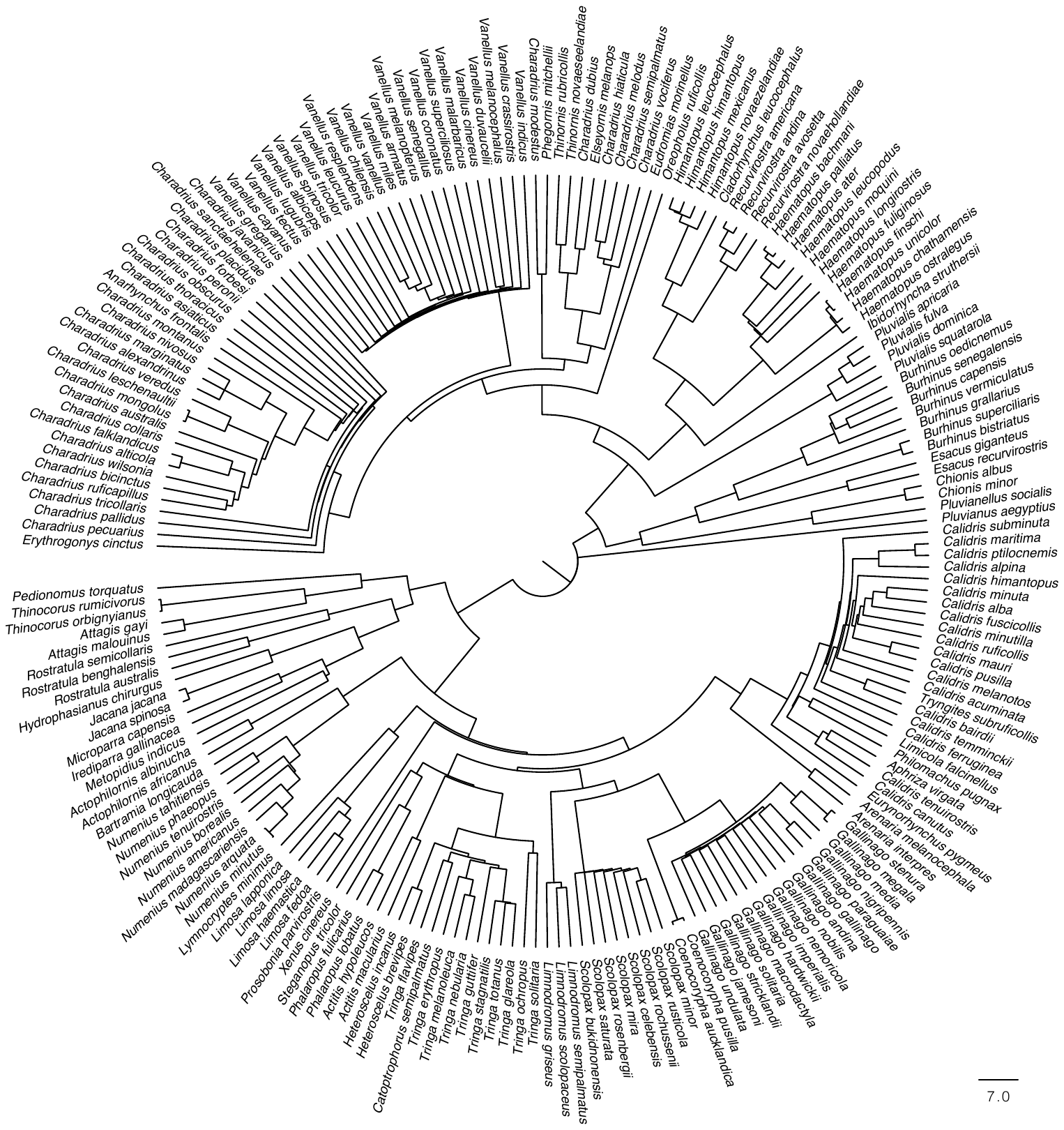

**FIGURE S1**. Phylogenetic hypothesis of 205 shorebird species used in this study.

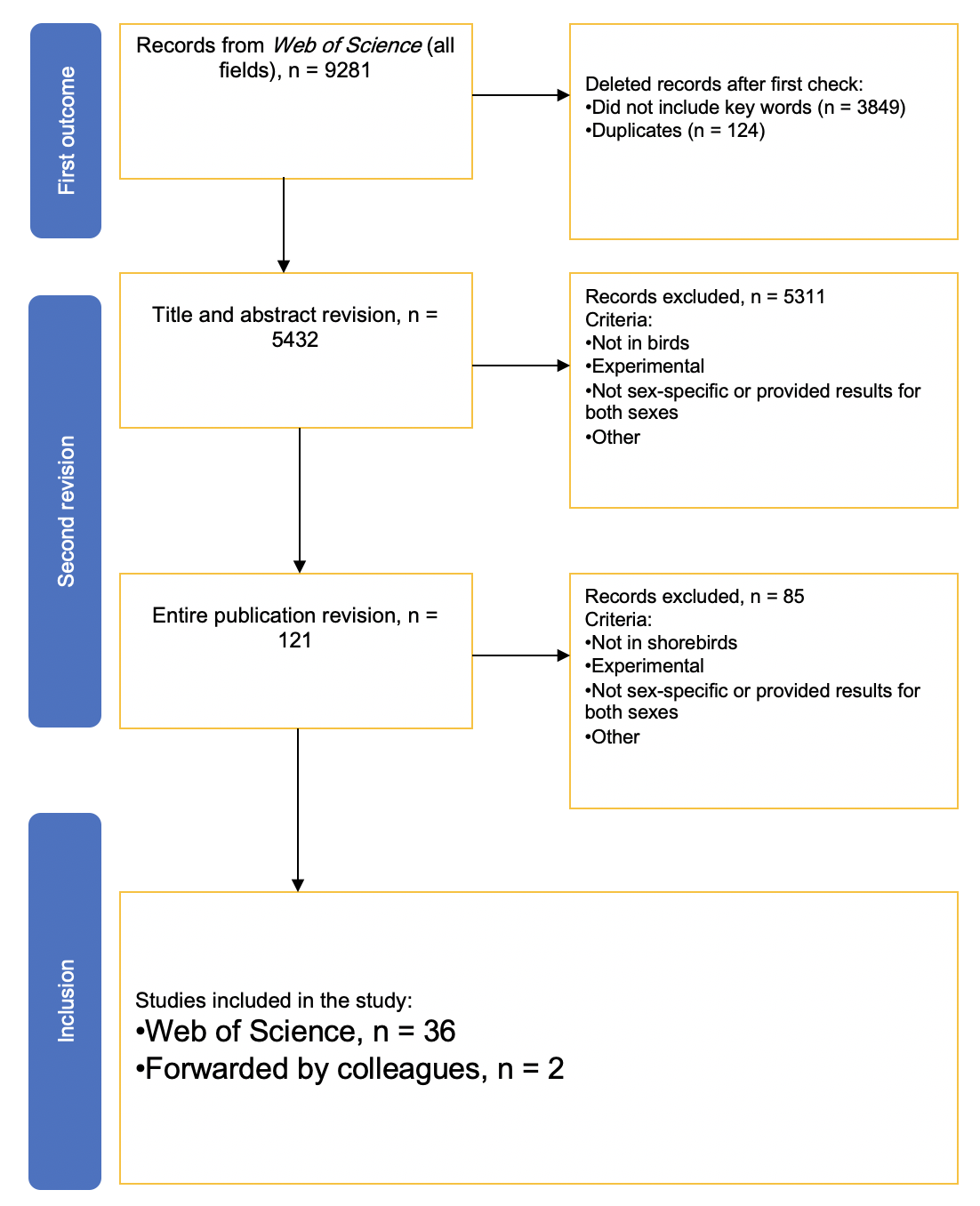

**FIGURE S2**. PRISMA Scheme and details of systematic literature search and inclusion.
